# PERK Kinase Activity is Regulated by Copper Binding-A New Regulatory Paradigm for Modulation of ER Stress Tolerance

**DOI:** 10.1101/2022.09.02.506340

**Authors:** Sarah E. Bond Newton, Noah R. Beratan, Cagla Akay-Espinoza, Margaret K. Bond, Xinglong Shi, Julia Perhacs, Tali Gidalevitz, Donita C. Brady, Kelly L. Jordan-Sciutto

## Abstract

PERK is a kinase that, in response to ER stress, mediates duplicitous homeostatic and pro-apoptotic signaling. Thus, intricate regulation is required for physiologic function. Attempts to modulate PERK activity have shown that the determinants of adaptive vs. mal-adaptive signaling remain ambiguous. Here, with purified protein, we provide direct evidence that PERK binds copper, identify the residues that coordinate binding, and demonstrate that binding is necessary for kinase activity. Furthermore cellular PERK activity can be modulated via copper availability, and this regulatory relationship can be manipulated to dictate ER stress tolerance. Critically, these phenomenon translate to phenotypes *in vivo*, as *C. elegans* harboring a “copper-binding mutant” of PERK exhibit enhanced ER-stress sensitivity. The copper-PERK paradigm adds onto a new class of dynamic copper-binding enzymes and implicates that copper homeostasis, as a regulator of PERK, may constitute a previously unknown variable to resolve long-standing ambiguity in endeavors to therapeutically target PERK.

## Introduction

Pancreatic/PKR-like endoplasmic reticulum (ER) kinase (PERK) is a kinase that phosphorylates the alpha subunit of eukaryotic initiation factor 2 (eIF2α) in response to the accumulation of unfolded proteins and other ER stressors^1,2^. In addition to PERK, there are three other eIF2α kinases that make up the integrated stress response (ISR). These kinases include protein kinase RNA-activated (PKR), general control nonderepressible 2 (GCN2), and heme-regulated inhibitor (HRI), which respond to specific cellular stresses, i.e., viral infection, amino acid deprivation, and heme insufficiency, respectively^3–5^. PERK responds to ER stress as part of a greater stress signaling network termed unfolded protein response (UPR), which aims to re-establish protein homeostasis^6,7^. Specifically, phosphorylation of eIF2α (p-eIF2α) causes global protein translation attenuation, with selective upregulation of transcripts encoding factors that mitigate stress, such as protein chaperones^1,2,8^. In addition, PERK phosphorylation of the transcription factor nuclear factor erythroid 2-related factor 2 (Nrf2) induces the antioxidant response^9,10^. Cumulatively, these signaling cascades underlie PERK’s role as a mediator of ER stress tolerance and survival, which has been demonstrated in numerous studies where ablation of PERK signaling enhances sensitivity to ER stress-induced cell death^4,8,11–20^. In complement, increasing the level or extending the duration of p-eIF2α signaling can mitigate ER stress-induced death^6,15,16^.

However, under prolonged or severe stress, subsequent signaling steps, prominently C/EBP homologous protein (CHOP) upregulation, result in the PERK-dependent induction of cell death to dispose of terminally stressed and damaged cells^21–24^. Given this biphasic nature, multiple feedback loops serve to tightly regulate PERK signaling. In the primary phase of adaptive signaling, calcium homeostasis networks and chaperones such as BiP and p58^IPK^ act both directly on PERK and indirectly through the mitigation of protein misfolding to curb signaling^25–28^. Furthermore, CHOP, a canonically pro-apoptotic transcription factor, also induces the expression of growth arrest and DNA damage-inducible protein 34 (GADD34), an adaptor protein which targets protein phosphatase 1 (PP1) to p-eIF2α to release the translational block^29^. This activity-dependent feedback loop, in conjunction with the constitutively expressed adaptor CReP^30^, provide the mechanism for oscillatory PERK signaling during unresolved stress, as observed experimentally and outlined in a mathematical model of UPR activity and outcomes as a factor of stress levels^31^.

Compellingly, utilizing that model, Erguler et al. define that low, intermediate, and high UPR activity states are associated with stress adaptation, tolerance, and initiation of apoptosis, respectively. While these responses, in turn, are linked to mild, moderate, and severe stress, respectively, they can also be reached in a number of different ways, depending on the history of stress and the difference in timescales used to define eIF2α phosphorylation/dephosphorylation versus the “inflicted genetic regulations,” such as BiP and CHOP accumulation. In particular, the authors noted that while intermediate activity as a result of translational block after high activity constitutes a commitment to apoptosis, intermediate activity following low activity that increases the folding capacity of the cell as part of stress adaptation constitutes tolerance through preconditioning. This preconditioning effect, also referred to as hormesis, has been documented in studies where preemptive induction of p-eIF2α enhances ER stress tolerance^15^. In general, hormesis refers to a form of stress adaptation, whereby exposure to a certain stress level acts as a preconditioning experience, preparing the cell for more severe stress in the future^32,33^. This is mediated by changes in the folding capacity of the cell, such as the induction of BiP and expansion of the ER, which outlast signaling effects, such as protein translation inhibition, and allow the resolution of equivalent stress when encountered a second time, with reduced/no PERK induction, and even the mounting of tolerance to subsequent greater stress with lower apoptotic commitment^15,34,35^. This PERK-dependent phenomenon is physiologically essential in tissues with high secretory capacity, and thus protein-folding loads, such as the pancreas where PERK is essential^36–38^.

In line with this multi-faceted, elastic feedback network as well as the context-dependent, disparate, and potentially severe nature of PERK signaling outcomes, PERK signaling is homeostatic in only a narrow, ill-defined window of activity in a goldilocks-type paradigm and requires precise regulation to achieve such, likely including factors not yet defined. Indeed, pathological dysregulation of UPR and ISR signaling, including PERK, has been extensively documented and proposed to play a causal role in many diverse pathologies characterized by chronic stress and aberrant cell fate outcomes^1,3,20,39^. However, pursuant attempts to target PERK signaling have led to mixed results.

For instance, in cancer, genetic and pharmacologic manipulations aimed at both the ablation and activation of PERK signaling have contrarily resulted in both disease exacerbation and mitigation depending on the model, intervention method, and measured outcome^40–43^. PERK’s pursuantly defined role as a “haploinsufficient tumor suppressor” is mechanistically understood as a combinatory effect. On one hand, stress tolerance and survival in the tumor microenvironment depends on adaptive PERK signaling; contrastingly, malignant transformation’s association with escape from cell cycle checkpoints and damaged cell disposal is mediated by, and therefore relies on, apoptotic PERK’s signaling. Despite such elucidations, the duplicitous implications have proved a detrimental limit in its effective therapeutic targeting thus far^44,45^.

On the other end of the spectrum, in response to the pathogenic protein misfolding in neurodegenerative disease, extensive evidence supports chronically elevated PERK-dependent eIF2alpha phosphorylation and CHOP signaling, ultimately aligning with the clinical presentation of neuron loss^21,34,46,47^. In conflict with the linear biphasic model, however, these chronic stress conditions still appear to utilize mitigating, if outbalanced, adaptive PERK signaling, as evidenced by contradicting lines of evidence^20,48^. For instance, risk for several neurodegenerative diseases has been linked to genetic variations associated with both hypo- and hyperactive PERK depending on the context, demonstrating the pathologic implications of UPR dysregulation, whether it results from impaired or excessive stress response^1,49–52^. In following, various interventions up- or down-regulating eIF2α phosphorylation, both in degree and duration, or its surrounding signaling components, have again proved to modulate neurodegenerative phenotypes in both therapeutic and exacerbating ends, depending on the model, suggesting that the adaptive and apoptotic functions of PERK signaling are simultaneously at play in these conditions, and again highlighting the shortcomings of the biphasic model^53–61^. Collectively, PERK’s spectrum of activity in disease pathology highlights the critical nature of maintaining its fine-tuned response throughout an organism’s life.

The oscillatory model resolves these still seemingly contradictory findings in a general manner, demonstrating that the dysregulated PERK signaling or accumulation of p-eIF2α in pathologic conditions can result not only from chronic or high PERK activity, as suggested by the biphasic system, but also from hypoactive PERK and impaired ER stress response pathways in the oscillatory model, where the degree to which this initial round of signaling resolves stress determines if one or more rounds of signaling will occur. This model highlights that it is the direction from which the oscillatory intermediate activity state is reached that determines apoptotic versus stress-tolerant outcomes. In other words, the activity of ISR and UPR over time, and not a particular threshold of eIF2α phosphorylation, dictates cell fate outcomes. Thus, effective therapeutic targeting of PERK requires a better understanding of other PERK modulators, such as those that may alter the difference in the timescales of eIF2α phosphorylation turnover and the resulting genetic regulations, as they define the relationship between stress and PERK activity levels, and thus the entry and outcomes of the critical intermediate state.

We propose copper, and its homeostasis in the cell, as such a modulator. Copper has vital physiologic roles, which have been classically attributed to its loading into essential proteins as a static cofactor, its redox cycling capabilities, and its toxicity in its free form^62,63^. However, growing evidence shows the presence of a pool of labile, lightly bound copper, which can be mobilized during signaling events to transiently affect protein function^64–66^. Turski et al. characterized the first copper-regulated kinases, mitogen activated kinase kinases 1 and 2 (MEK1/2)^67–70^. Thereafter, several other kinases and other enzymes, including Unc-51 like autophagy activating kinases 1 and 2 (ULK1/2), pyruvate dehydrogenase kinase 1 (PDK1), casein kinase 2 (CK2), phosphodiesterase 3B (PDE3B), a clade of ubiquitin-conjugating enzymes (UBE2D), and the histone H3-H4 tetramer have been characterized to bind and be regulated by copper^65,71–75^. The autophagic kinase ULK1/2 was specifically investigated in studies following the identification of MEK1/2 to identify the copper kinome using a tri-faceted screen: 1) mass spectrometry identification of kinases pulled down from cell lysates using copper-charged resin; 2) inhibition with the copper chelator tetrathiolmolybdate (TTM) in a high-throughput *in vitro* kinase assay; 3) sequence homology with the putative copper-binding sites identified in MEK1/2. PERK was another potential copper-regulated kinase identified by all three of these screens. Here, we validate that PERK is regulated by copper binding and provide evidence that copper homeostasis is a previously unappreciated variable in defining the outcomes of PERK signaling, namely ER stress tolerance.

## Results

### PERK binds copper near the kinase active site

Given the aforementioned preliminary screen results suggesting PERK’s ability to interact with copper, PERK’s copper-binding activity was first validated via its ability be pulled down from mouse embryonic fibroblast (MEF) whole-cell lysates using immobilized metal affinity chromatography (IMAC) followed by western blotting. As shown in **Figure 1A**, PERK was selectively pulled down by copper-charged IMAC resin, as opposed to uncharged or zinc-charged resin. Furthermore, other abundant cellular proteins, such as actin, were not pulled down (**Figure 1A**). To determine whether this interaction was direct or required cellular mediators, the interaction of purified recombinant GST-tagged PERK kinase domain with differently charged IMAC columns was also assessed. As shown in **Figure 1B**, purified PERK was again selectively pulled down by the copper-loaded column, though some interaction was also seen with zinc-charged resin (**Supplemental Figure 1A**).

**Figure 1:**
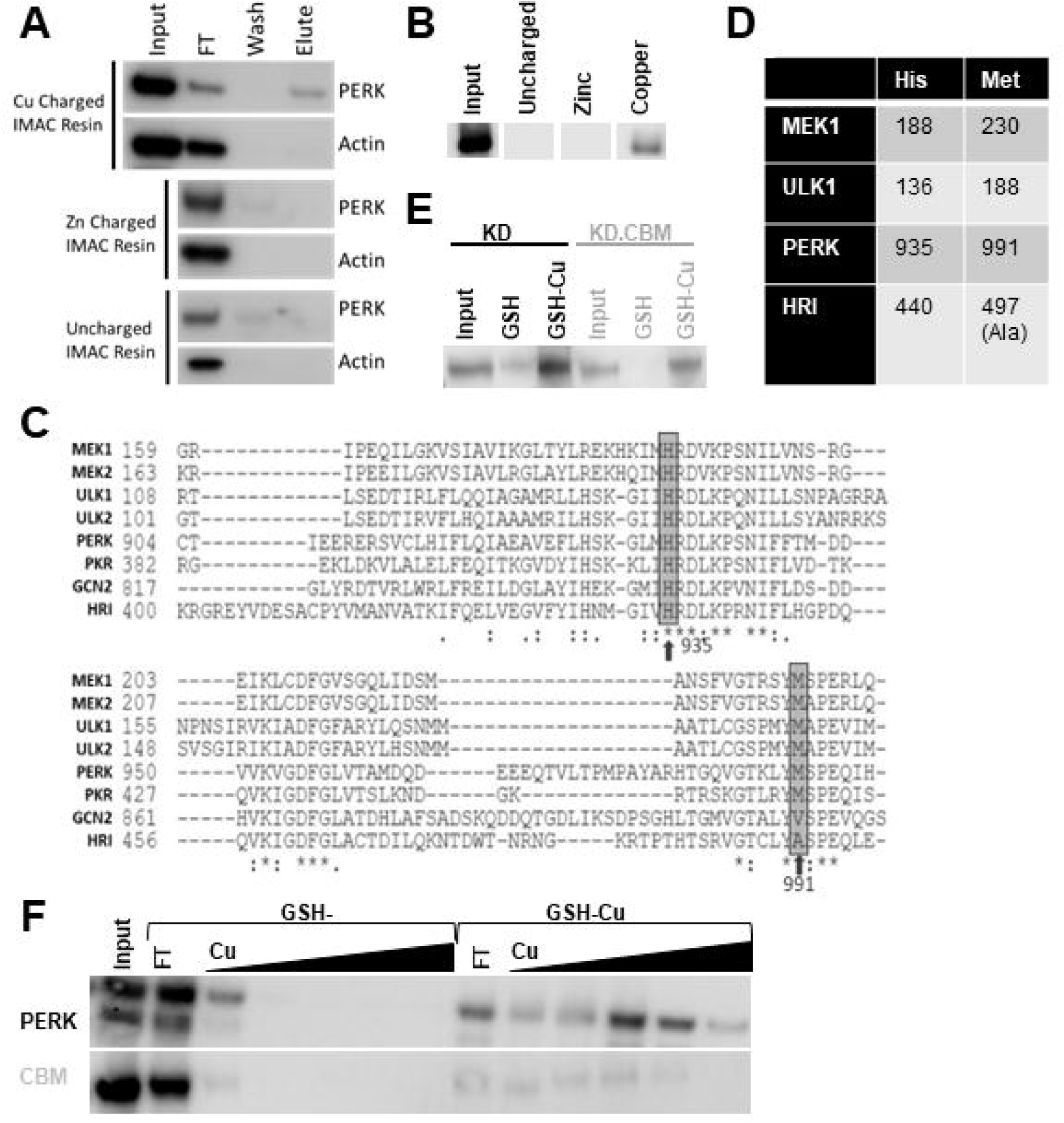
PERK binds copper near the kinase active site. **A,B)** Western blot images, probed for PERK or as indicated, of 100µg of MEF whole cell lysate(A) or 100ng of purified GST-PERK kinase domain protein(B) pulled-down by IMAC resin, charged as indicated, and eluted with imidazole. FT=Flow-through. Wash=Binding buffer+20mM imidazole. Elute=Binding buffer+100mM imidazole. **C)** Alignment of PERK amino acid sequence with other eIF2α kinases and previously characterized copper binding proteins MEK1/2 and ULK1/2. Putative copper binding residues are outlined, with the residue numbers for PERK indicated. **D)** Table dictating putative copper binding residues by homology. **E,F)** Western blot images, probed for PERK, of 500ng of purified WT or CBM (gray) kinase domain(KD, E) or full-length(F) PERK-FLAG proteins pulled-down by glutathione sepharose, uncharged or charged with copper, and remaining bound after washes (E) or eluted with copper sulfate (F, 33µM-33mM, complexed 1:3 with glutathione). Also see Supplemental Figure 1.

To determine the residues coordinating copper binding, PERK’s putative copper-binding site was identified based on alignment with previously characterized copper-binding proteins (**Figure 1C-D**). Specifically, histidine 935 and methionine 991, located near the active site in PERK’s kinase domain, were mutated to alanine to form the copper-binding mutant (CBM). The copper binding activity of wild-type (WT) and CBM constructs was assessed using glutathione sepharose, which has been shown to be able to bind copper and hold it in its reduced form^63,76–78^. The purified PERK kinase domain was pulled down by uncharged glutathione sepharose, while the CBM protein showed no interaction, suggesting the presence of copper in the purified WT but not the CBM protein (**Figure 1E, Supplemental Figure 1B**). On the other hand, both proteins were able to interact with sepharose which was pre-charged with copper, suggesting that the mutation of the copper-binding domain resulted in reduced, rather than the completely ablated, copper-binding activity, such that interaction with copper could be facilitated by locally high concentrations of the copper-charged resin (“GSH-Cu”, **Figure 1 E, Supplemental Figure 1B**). Similar interactions were observed with the purified full-length WT and CBM PERK proteins, although competitive elution with copper sulfate suggested that the WT protein had higher affinity for the copper-charged resin (**Figure 1F, Supplemental Figure 1C**). These results indicate the PERK is able to bind copper directly both in the cellular environment and *in vitro* and that histidine 935 and methionine 991 together coordinate this binding.

### Copper is required for PERK kinase activity in vitro

After determining PERK’s copper-binding activity, we next sought to test whether this binding regulated kinase activity using an *in vitro* kinase assay assessed by both ELISA and western blotting. Indeed, as shown in **Figure 2A-B**, the full-length CBM PERK was kinase-dead in comparison to the full-length WT PERK, even at time points well beyond the linear phase on the enzyme progress curve, when product accumulation was observed to plateau for the WT protein (**Supplemental Figure 2A-E**).

**Figure 2:**
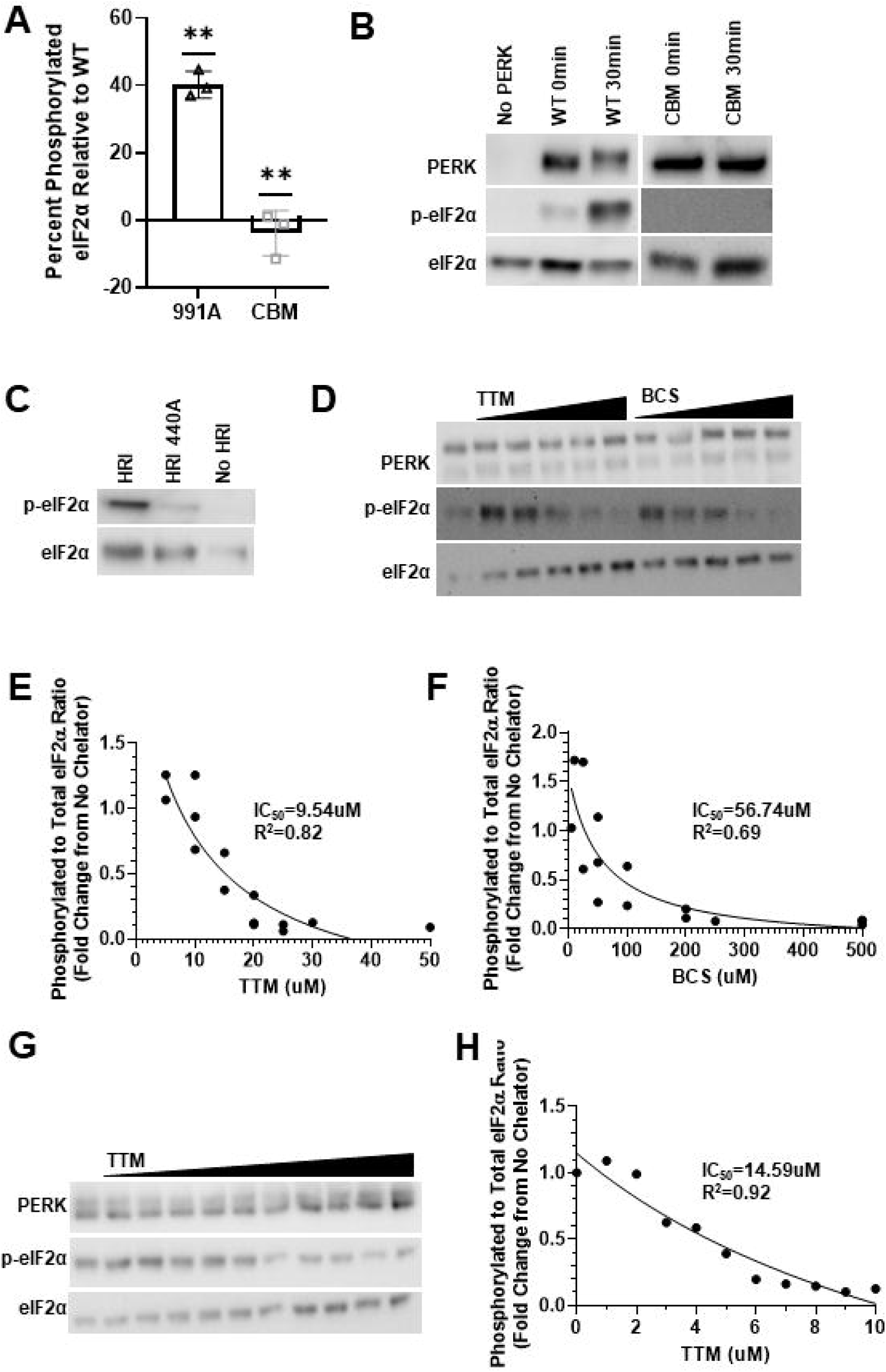
Copper is necessary and sufficient for PERK-eIF2*α* phosphorylation in the cell. **A)** Quantification of p-eIF2α levels from full-length PERK-FLAG protein kinase reactions carried out at 37°C, normalized to WT levels at that time point (approximately 7 minutes) as monitored by ELISA absorbance at 450nm. Quantification plots individual data points from experimental replicates, n=3, and each bar represents mean±SD. One-sample t-tests with a theoretical mean of 100 (WT) returned **p<0.01 for 991A (black triangles) and CBM (gray squares). A two-tailed paired t-test for 991A vs CBM also gave p<0.05. **B,C)** Representative western blot images, probed as indicated, of full-length PERK-FLAG(B) or HRI-FLAG(C) kinase assay reactions carried out at 37°C for indicated time points(B) or one hour(C). **D,G)** Representative western blot images, probed as indicated of full-length WT PERK-FLAG(D) or GST-PERK kinase domain(G) kinase assay reactions carried out at 37°C for 10(D) or 5(G) minutes in the presence of increasing titrations of copper chelators TTM (10-30µM, B), 1-10µM, G), or BCS(25-500µM). **E, F, G)** Quantification of n=3 variable replicates of the experiment shown in D(E,F) or G(H, n=1); data fit by least squares regression to a three-parameter dose-response inhibition curve. IC_50_ and R^2^ of the fit curves indicated. Also see Supplemental Figure 2.

One of the two residues mutated in the CBM, histidine 935, is part of the HRD motif, which is well conserved among kinases, and thus may be important for kinase activity in a manner independent of copper-binding activity. Therefore, we assessed whether the kinase-dead CBM PERK was a result of impaired copper-binding ability. To address this, we tested the effect of analogous mutations in another eIF2alpha kinase, HRI, that was not identified in the initial screen for copper-binding kinases. As shown in **Figure 1C-D**, HRI, which retains an aligned HRD histidine at residue 440, has an alanine at residue 497, compared with PERK’s methionine at 991. As such, a single mutation resulting in an alanine at residue 440 in HRI created an analogous change to those introduced into the kinase domain of CBM PERK. As expected, this mutant HRI showed diminished kinase activity relative to the unmutated HRI but lacked the kinase-dead phenotype of CBM PERK **(Figure 2C, Supplemental Figure 2F-G**), confirming that this histidine in the HRD motif was not essential for kinase activity, even in conjunction with the presence of alanine at the second residue, and suggesting functionality, i.e., binding copper,for this pair of residues that is not universal among kinases. Furthermore, a single mutation at methionine 991 to alanine (991A), which appears to be conserved only in copper-regulated kinases (**Figure 1C-D**), resulted in a protein which exhibited partial kinase activity significantly different from both the WT and CBM PERK proteins (**Figure 2A, Supplemental Figure 2A-C**), again supporting the specific functionality of these residues for copper-regulated kinase activity and suggesting a dose-response relationship between PERK-copper binding and PERK kinase activity.

We employed additional measures to confirm that PERK-copper binding regulated PERK in a dose-dependent manner. Two copper specific chelators, bathocuproine disulfonic acid (BCS) and TTM, inhibited *in vitro* PERK kinase activity, such that p-eIF2α levels could be well fitted to an inhibition curve against increasing concentrations of the chelator (**Figure 2D-F, Supplemental Figure 2D-E**). Similar trends were observed with the GST-tagged PERK kinase domain and TTM (**Figure 2G-H, Supplemental Figure 2H-I**). Overall, these results indicate that PERK requires copper binding for kinase activity and that copper binding can be used to modulate PERK kinase activity in a dose-dependent fashion.

### Copper is necessary and sufficient for PERK-eIF2***α*** phosphorylation in the cell

We next sought to translate our *in vitro* findings into the cellular context by determining whether manipulating intracellular copper levels affected physiologic PERK activity. MEFs were pretreated with the copper chelators BCS and TTM, followed by treatment with pharmacological PERK inducers, PERK activator (PA) and thapsigargin (Tg). In these experiments, CCS, the stability of which has an inverse relationship with copper, was used as an indicator for effective copper chelation (**Figure 3A,C, Supplemental Figure 3A,C**)^79,80^. As shown in **Figure 3A-B**, BCS significantly inhibited eIF2α phosphorylation in response to treatment with PA. Similarly, TTM significantly inhibited eIF2α phosphorylation in response to treatment with Tg, a canonical ER stress inducer (**Supplemental Figure 3A-B**).

**Figure 3:**
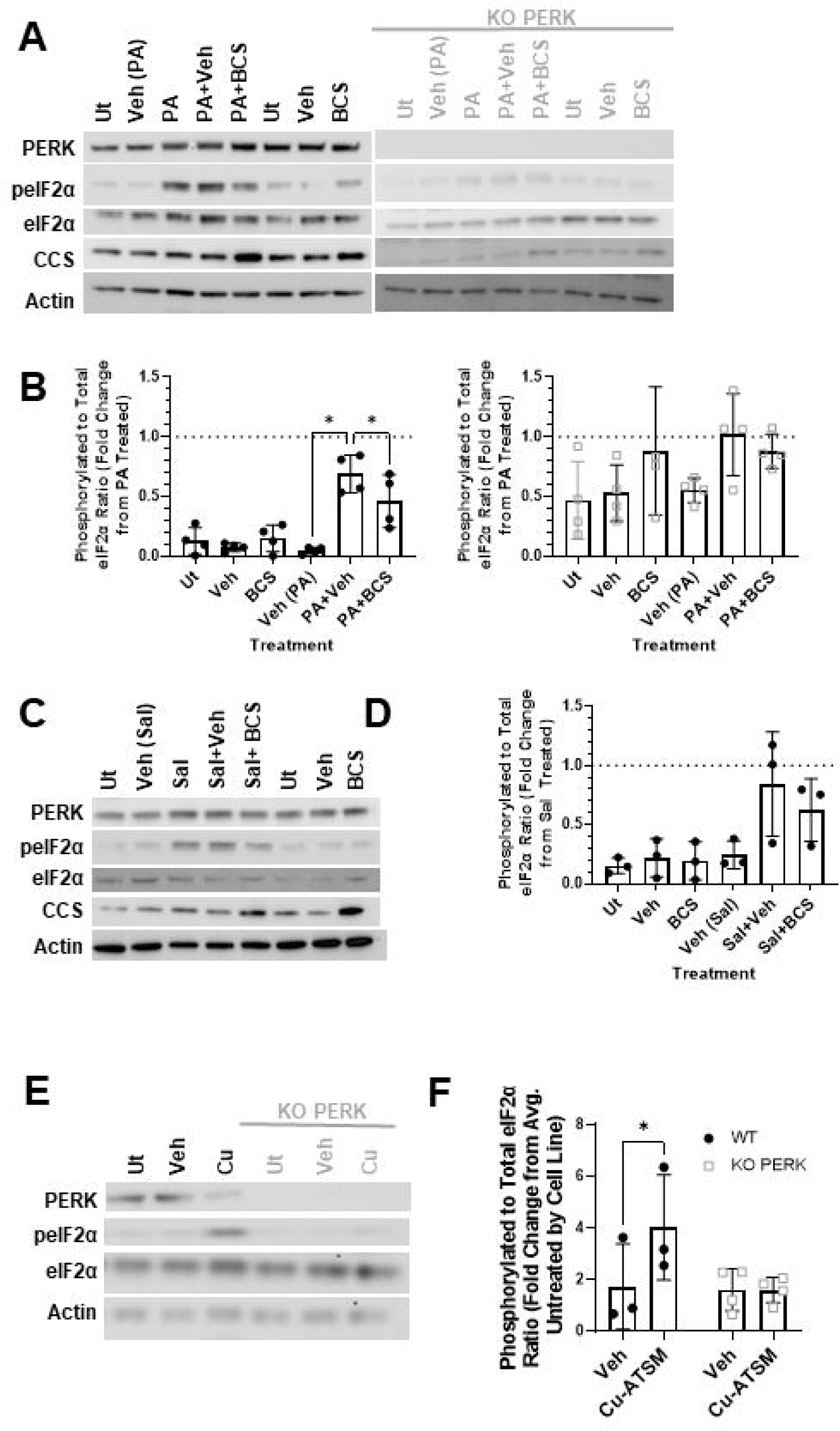
Copper is necessary and sufficient for PERK-eIF2*α* phosphorylation in the cell. **A,C,E)** Representative western blot images, probed as indicated, of WT and KO PERK (gray) MEF whole cell lysates from cells treated with 500µM BCS for 24 hours, and 10µM PA for 1 hour(A) or 50µM Sal for 24 hours(C), or 100µM Cu-ATSM alone for 1 hour(E). **B,D)** Quantification of n=3-4 replicates of the experiments in A(B) or C(D). Data is normalized by experimental replicate as fold changes from p-eIF2α induction alone(PA or Sal) as indicated by the dotted line at y=1. One-way repeated measures ANOVA returned no significant effect for Sal treatment (D, p=0.054) or PA treatment of KO PERK MEFs (B, gray squares, p=0.163), but returned a significant effect for WT (p=0.0046), with Holm-Sidak’s multiple comparison tests returning significance for each group in comparison to PA+Veh: *p<0.05, **p<0.01. **F)** Quantification of n=3-4 replicates the experiment in E. Data is normalized as the fold change from the average untreated ratio by cell line as indicated. Two-way repeated measures ANOVA returned treatment (p=0.0415) and treatment/genotype interaction (p=0.0223) as significant factors. Sidak’s tests for multiple comparisons between treatments by genotype returned *p<0.05 for WT. All quantifications plot individual n as data points and each bar represents mean±SD. Also see Supplemental Figure 3.

In PERK knockout MEFs, however, neither Tg nor PA significantly induced eIF2α phosphorylation and, in following, chelation of copper had no significant effect (**Figure 3A-B, Supplemental Figure 3A-B**). When salubrinal (Sal), an inhibitor of eIF2α dephosphorylation, was used to cause accumulation of phosphorylated eIF2α in a non-PERK-dependent fashion, neither chelator had a significant effect, although salubrinal also failed to significantly increase eIF2α phosphorylation (**Figure 3C-D, Supplemental Figure 3C-D**). The common trends observed across these cell types and pharmacological manipulations, given their different mechanisms of action, suggest a threshold of copper availability in the cell as a requirement for PERK induction. Furthermore, treatment with the copper ionophore Cu-ATSM was sufficient to induce eIF2α phosphorylation in a PERK-dependent manner in the absence of stress (**Figure 3E-F**), suggesting that copper is not only necessary but also sufficient for PERK induction.

### Copper homeostasis dictates PERK signaling and ER stress tolerance

Given the ability of changes in intracellular copper availability to specifically modulate PERK as presented in Figure 3, we sought to validate a hypothetical model where maintenance of copper homeostasis is required for precise PERK regulation and thus physiologic ER stress tolerance. CTR1 is the cell’s main copper importer, and knockout cells without CTR1 (KO CTR1) should have lower copper content^81^. Therefore, we compared response to Tg in MEFs isolated from littermate CTR1 and KO CTR1 mice. KO CTR1 MEFs had a significantly diminished response to Tg in comparison to CTR1 MEFs, after adjusting for baseline differences in p-eIF2α, which were also significantly different (**Figure 4A-B, Supplemental Figure 4A**), indicating that copper-depleted cells had a stressed basal state and impaired PERK-eIF2α phosphorylation following ER stress. Notably, these findings were reminiscent of KO PERK MEFs, as observed here and in previous investigations (**Figure 3A, Supplemental Figure 3A**)^4,22,36,44^. In following, as previously reported, KO PERK MEFs had impaired ER stress tolerance, specifically greater cytotoxicity, measured by lactate dehydrogenase (LDH) release, in response to 20 hours of ER stress induction by Tg or tunicamycin (Tm) (**Supplemental Figure 4C-D**)^19^. In KO PERK MEFs, the impaired ER stress tolerance was complemented by the accumulation of p-eIF2α, indicating prolonged, unresolved stress, as well as the accumulation of cleaved caspase 3 independent of CHOP induction in comparison to WT MEFs (**Supplemental Figure 4E**). Unexpectedly, no corresponding significant differences in cytotoxicity were observed between the CTR1 and KO CTR1 MEFs in response to Tg or Tm treatment, whereas p-eIF2α, CHOP, and cleaved caspase 3 accumulation were observed in both lines (**Figure 4C-E**).

**Figure 4:**
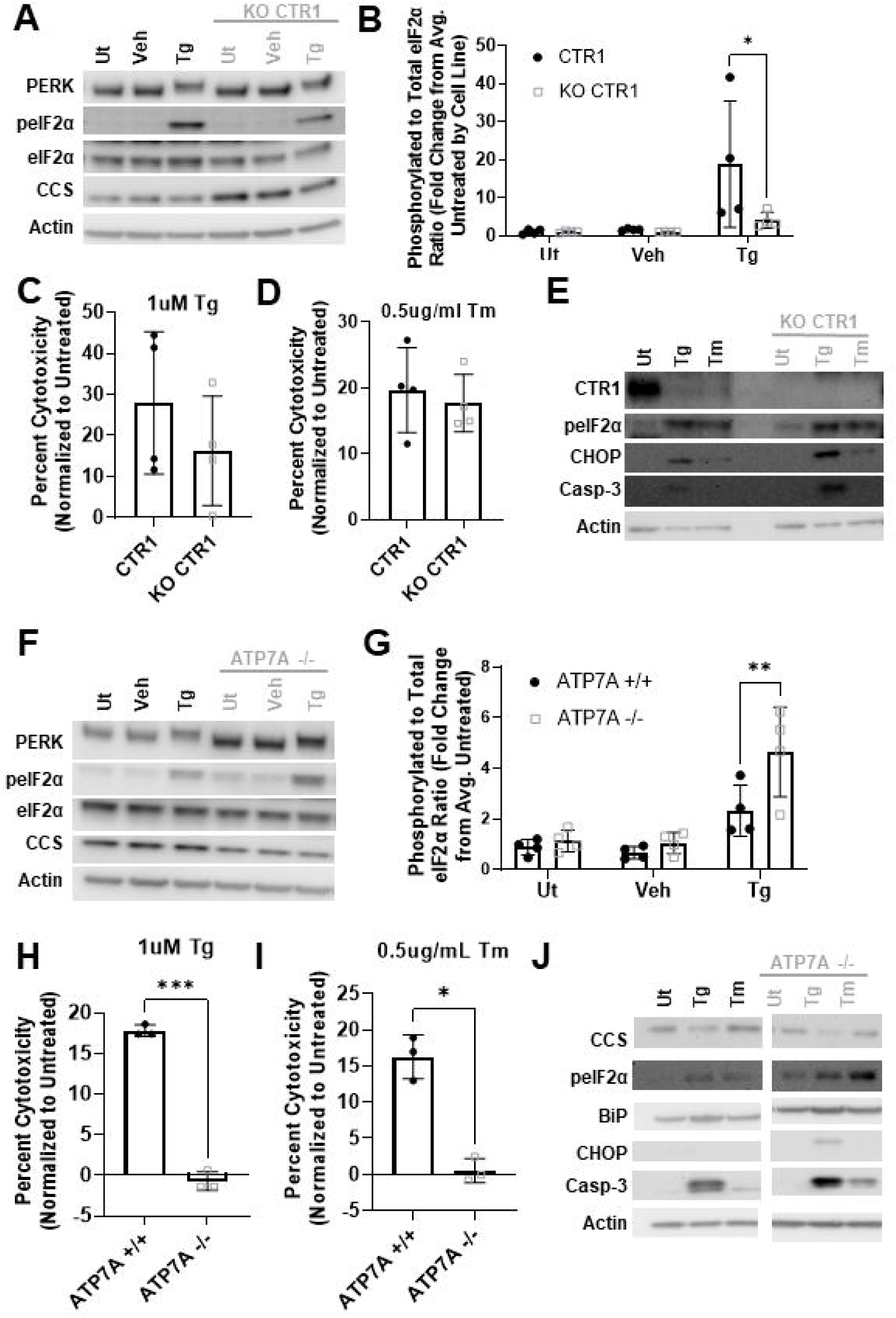
Copper homeostasis dictates PERK signaling and ER stress tolerance. **A,E,F,J)** Representative western blot images, probed as indicated, of CTR1 and KO CTR1 (gray)(A,E) or ATP7A +/+ and ATP7A −/−(gray) (F,J) MEF whole cell lysates from cells treated with 300nM Tg for 2 hours (A,F), or 300nM Tg or 0.5ug/mL Tm for 20 hours (E,J). E and J images are representative of n=3 biological replicates. **B,G)** Quantification of n=4 replicates of the experiments in A(B) or F(G). Data is normalized as the fold change from the average untreated ratio, pooled or by cell line as indicated. Two-way repeated measures ANOVA returned treatment as a significant factor (p=0.0367) for the CTR1 comparisons(B) and both treatment (p=0.0012) and genotype (p=0.0053), as well as their interaction (p=0.0212) as significant factors for the ATP7A comparisons(G). Sidak’s tests for multiple comparisons each treatment between genotypes returned significance for Tg in both pairs: *p<0.05; **p<0.01. **C,D,H,I)** LDH Assay results quantifying LDH release as a measure of cytotoxicity for CTR1 and KO CTR1 (gray squares)(C,D), or ATP7A +/+ or ATP7A −/− (gray squares)(H,I), MEFs treated with 1µM Tg(C,H) or 0.5ug/mL Tm(D,I) for 20 hours. Cytotoxicity is calculated as a percentage of the total LDH released by lysed cells, after normalization by subtracting out any LDH release signal from untreated cells. Two-tailed paired t-tests: *p<0.05; ***p<0.001. All quantifications plot individual n as data points (n=3-4) and each bar represents mean±SD. Also see Supplemental Figure 4.

In parallel, we examined response to Tg in a pair of MEF lines with the ATP7A gene either expressed (ATP7A +/+) or not (ATP7A −/−). Given that ATP7A is a copper exporter, its deletion resulted in higher intracellular copper content, as suggested by lower CCS levels (**Figure 4F**)^81^. While baseline eIF2α phosphorylation was not different between ATP7A −/− and ATP7A +/+ cells (**Supplemental Figure 4B**), significantly higher eIF2α phosphorylation was observed upon Tg treatment in ATP7A −/− cells (**Figure 4F-G**), demonstrating that acute PERK activity in response to ER stress may be bidirectionally modulated by intracellular copper availability and that, consequently copper may be an endogenous regulator of PERK signaling. Indeed, as shown in **Figure 4H-I**, nearly no release of lactate dehydrogenase was observed in ATP7A −/− MEFs in response to ER stress inducers that resulted in observable cytotoxicity in ATP7A +/+ MEFs, suggesting that these copper-replete cells were resistant to ER stress, even though both cell lines exhibited notable accumulation of p-eIF2α and cleaved caspase 3 during long-term stress (**Figure 4J**). These findings demonstrate that copper itself can be manipulated to modulate PERK signaling and pursuant ER stress tolerance.

### In vivo ER stress tolerance is impaired in the presence of CBM PERK

Given the above evidence for the molecular regulation of PERK by copper binding and the implication of a role for copper homeostasis in ER stress tolerance as shown in Figure 4, we next sought to determine if these findings translated *in vivo* utilizing an animal model. Alignment of the amino acid sequences of PERK homologues across various model systems showed that the identified copper-binding residues were conserved among multiple species including *Caenorhabditis elegans* at residues histidine 931 and methionine 991 (**Figure 5A**). Many aspects of the UPR, particularly PERK signaling, have been dissected in *C. elegans* in clear phenotypic assays, making it a compelling model for the investigation of PERK-dependent physiologic phenotypes^82–87^. Unique PERK-dependent hormetic effects have also been reported in *C. elegans*^88^. Most notably, *C. elegans* with KO pek-1, the PERK homolog, while albeit generally normal in phenotype, have been reported to exhibit increased reliance on the ire-1 signaling arm of the UPR for development and to have dramatically impaired ER stress tolerance^89–91^. Specifically, RNAi knockdown of xbp-1 in pek-1 KO *C. elegans* results in developmental arrest prior to the L2 larval stage and Tm treatment results in a higher percentage of death and maturation arrest compared to the WT controls^89^. Thus, we used *C. elegans* to test the physiologic relevance of PERK-copper binding and regulation at the organism level.

**Figure 5:**
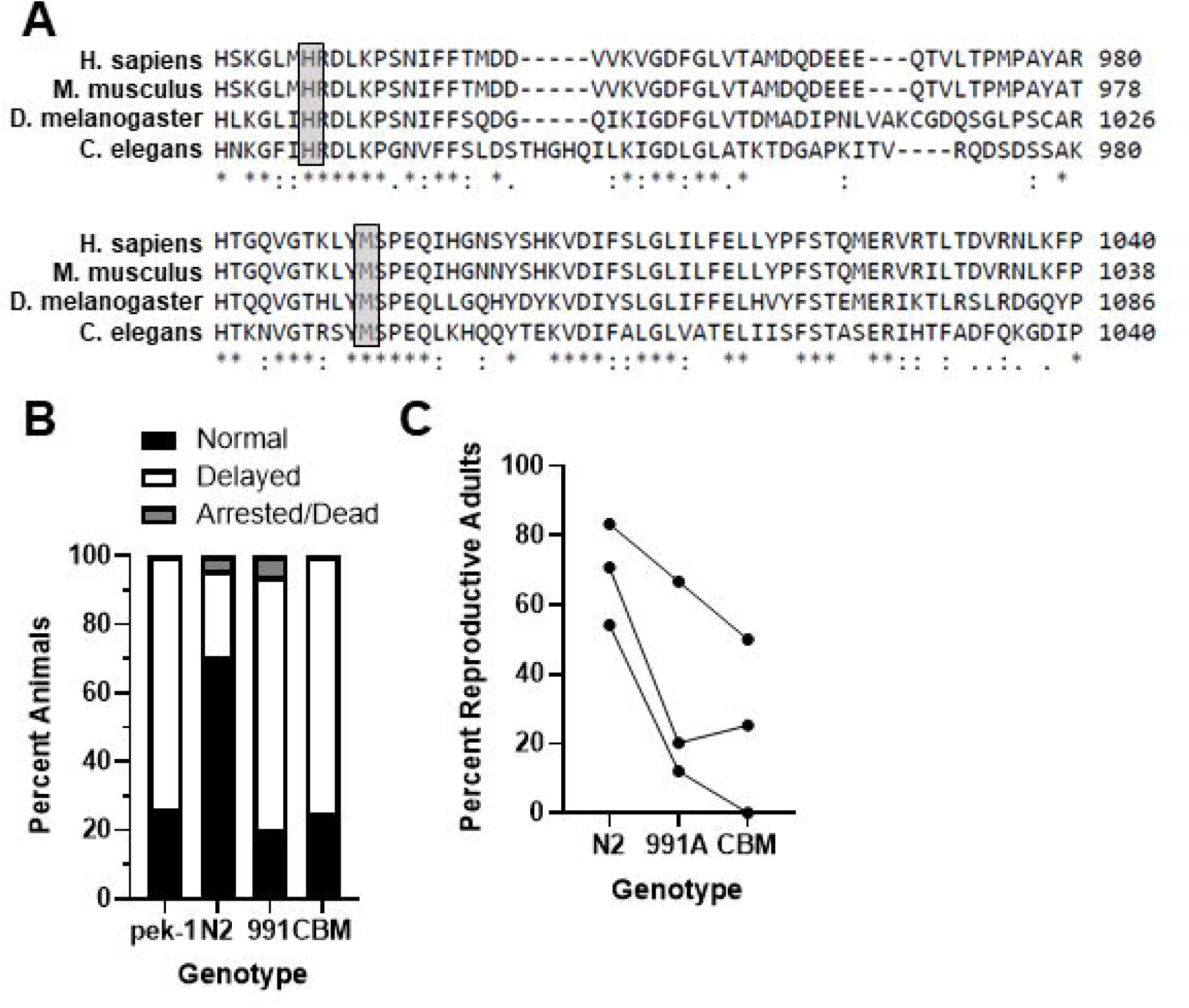
Copper-binding mutant PERK animals have impaired ER stress tolerance. **A)** Alignment of PERK amino acid sequence showing conservation of binding residues (highlighted) across organisms. **B)** Development/Survival analysis of *C. elegans* eggs of each strain incubated on 1ug/mL Tm for 3 days. Chi-squared=73.03, p<0.0001. **C)** Plot of percentage of *C. elegans* developed to reproductive age by day 3 in 3 biological replicates of the experiment in 5B. One-way repeated measures ANOVA returned a significant effect for genotype (p=0.0098), with a left-to-right multiple comparison test for linear trend returning p=0.0048 and a slope estimate of −22.17 with a standard error of −3.931. Also see Supplemental Figure 5.

Using CRISPR editing, *C. elegans* were mutated to harbor alanine at the 991 copper-coordinating methionine, followed by at histidine 931 to create two *pek-1* mutant strains, *knu1130* (991) and *knu1138* (CBM). While these mutants exhibited no overt phenotypes under standard strain maintenance conditions, they were completely unable to develop past larval stage 2 when *ire-1* was knocked down by RNAi, similar to what was observed in pek-1 KO *C. elegans*, while the WT N2 *C. elegans* progressed through development normally. Furthermore, incubation on plates containing Tm resulted in death and developmental delays in the CBM *C. elegans* compared to the WT N2 *C. elegans*, again similar to those observed in pek-1 KO *C. elegans* (**Figure 5B-C, Supplemental Figure 5A-B**). Intriguingly, the putatively partially active single mutant (991A) also exhibited impaired ER stress tolerance, the level of which was between that observed in the WT and CBM phenotypes across replicates (**Figure 5C**). No significant differences were observed between the *C. elegans* lines under vehicle conditions (**Supplemental Figure 5C**).

## Discussion

In this study, we demonstrate that 1) PERK binds copper and identify the coordinating residues (**Figure 1**); 2) PERK-copper binding is required for PERK kinase activity (**Figure 2**); and 3) this interaction is necessary and sufficient for PERK induction in cells (**Figure 3**). We also demonstrate that the manipulation of copper homeostasis can modulate ER stress tolerance at the cellular level via PERK activity (**Figure 4**), and that this novel PERK-copper regulation paradigm has phenotypic *in vivo* significance at the organism level (**Figure 5**). Altogether, these results demonstrate that copper binding is a physiologic modulator of PERK activity and have broad implications for PERK signaling, its regulation, and its downstream effects on proteostasis and cell fate outcomes during ER stress in both physiology and disease.

In Figure 1, we showed that PERK interacts with copper in both cellular lysate and purified conditions. *In vitro*, PERK has some ability to bind zinc, although this interaction was weak, indicated by its ability to be perturbed by a non-competitive wash, suggesting that this observation may be an artifact of the *in vitro* environment and not a specific interaction. However, dual-specific zinc- and copper-binding pockets have been characterized in other proteins, such as SOD1, and this study does not rule out this possibility^62,63^. We identified histidine 935 and methionine 991 as two residues that play a necessary role in PERK’s regulation by copper and confirm that mutation of these residues impairs copper binding. The ability to pull down PERK with uncharged glutathione sepharose was unexpected; however, given the well-documented copper-binding activity of glutathione^77,78^, this finding suggests that WT PERK is at least partially loaded with copper under the experimental conditions for purification used here, whereas CBM PERK, presumably due to lower copper affinity, is not purified with the copper-binding site occupied and is thus unable to be captured by the uncharged resin. The copper-charged resin represents a condition of locally high copper concentrations, and, as such, its ability to pull down both constructs demonstrates that the CBM has a reduced, rather than ablated, copper affinity.

In Figure 2, we demonstrated that copper is required for PERK kinase activity, as the mutation of the binding residues resulted in diminished activity individually and a kinase-dead phenotype in combination. Intriguingly, the residues responsible for these phenotypic changes have also been noted to be mutated in tumors from The Cancer Genome Atlas and, although the effect and contribution of these mutations have not been directly evaluated, in general, hypomorphic PERK mutations have been noted across cancers, in agreement with the effects found in this study^44,92,93^. The ability to dose-dependently inhibit kinase activity with copper chelators provides further evidence that PERK retains copper when purified under these conditions, as discussed above. Lacking a structure of PERK with copper bound, the mechanism of copper-mediated PERK activity remains purely hypothetical; however, several possibilities in agreement with available literature are worthy of note. The utilization of the histidine in the HRD motif for coordination of copper as well as the absolute requirement of copper for kinase activity, be it auto-phosphorylation, as suggested by the lack of a molecular weight shift in CBM PERK, or substrate phosphorylation, suggest that copper has a direct catalytic activity on the kinase reaction, plausibly via stabilization or coordination of the flexible catalytic loop^69,94^.

In Figures 3 and 4, we provided evidence that cellular copper availability can be manipulated to control acute PERK activity and, as shown in Figure 4, we further determined whether such modulation constituted a mechanism to regulate ER stress tolerance during prolonged stress. While PERK signaling is canonically required for acute ER stress tolerance and has been proposed as a dictator of cell fate outcomes in various physiologic and pathologic conditions involving chronic ER stress, the literature has proven that the determinates of physiologic or adaptive activities versus pathologic or aberrant signaling remain ambiguous^1,3,4,45^. The findings here suggest that the evaluation of copper homeostasis in disease states may offer insights into whether and to what end targeting PERK may be therapeutic and further suggest that the manipulation of cellular copper availability itself may be a tool to achieve this goal.

We hypothesized that the heightened signaling observed with acute Tg treatment in copper-replete ATP7A−/− cells would result in improved ER stress tolerance with prolonged treatment, based on studies demonstrating that the enhancement or prolongation of p-eIF2α signaling mitigates ER stress-induced death^15,16^. Indeed, we observed not merely attenuation but the complete ablation of cytotoxicity in ATP7A−/− cells treated long-term with Tg or Tm, compared to the approximately 15% cytotoxicity observed in ATP7A +/+ cells under the same conditions. Remarkably, this was observed across a log range of concentrations for these stressors (data not shown). Conflictingly, the signaling pathways observed in these cells are similar to all other cell lines in which cytotoxicity is observed, including the accumulation of p-eIF2α, CHOP, and cleaved caspase 3. Recalling the mathematical model of UPR activity, we propose that these phenotypes reflect a stress-tolerant “senescent” state described by the authors, where commitment to apoptosis is avoided^31^. Given that the cell-fate dictating, key oscillatory intermediate activity state described in the model is a function of the difference in the time scales of PERK signals, we further propose that the difference is greater in ATP7A −/− cells due to a shortened eIF2α phosphorylation/dephosphorylation turnover time mediated by higher copper occupancy of PERK at the basal state, which results in quicker and higher PERK activation upon stress. In our working model, this “priming by occupancy” differs from preconditioning, as discussed for KO CTR1 cells below, because under basal conditions negative regulatory factors such as BiP keep PERK inactive even when it is occupied by copper. Thus, the unique phenotype of complete ER stress resistance, rather than enhanced tolerance, is supplied by a heightened initial peak activity and altered signaling kinetics, rather than by a greater folding capacity mediated by previous rounds of PERK signaling, which still results in apoptosis when exceeded.

In KO CTR1 MEFs, given their expected and demonstrated diminished eIF2α phosphorylation in response to short-term stress, we hypothesized that prolonged treatment would result in enhanced sensitivity to ER stress-induced death, relative to control, similar to that observed in KO PERK MEFs, as previously reported and recapitulated here^19,22,36^. Instead, we did not find differences in cytotoxicity, which may be a result of the inhibition, rather than the ablation, of PERK activity in KO CTR1 cells, which still mount a significant p-eIF2α response to Tg in the short term. Thus, while the initial PERK response is stunted, the accumulation of unresolved stress over multiple PERK signaling oscillations before that monitored at 20 hours may induce sufficient homeostatic signaling to maintain survival. Furthermore, the significantly different p-eIF2α levels in KO CTR1 MEFs compared to CTR1 MEFs under basal conditions suggest that hormesis may also play a role. Copper deficiency may impair PERK activity cycles required for basal cell survival and replication, leading to a stressed basal state. Due to this constant disruption of copper homeostasis, analogous to the sustained folding stress in pancreatic cells where PERK was initially identified, these cells may have engaged compensatory mechanisms to enhance folding capacity^38^. This stress-adapted baseline, in combination with the impaired magnitude of initial stress response, is reminiscent of the mathematical UPR model where entry into an intermediate activity state from a low, rather than a high, activity state results in stress tolerance, which is characterized by oscillations in translational attenuation and apoptotic signaling.

In CTR1 MEFs, we also observed a dramatic down-regulation of CTR1 after 20 hours of stress induction. Based on previous literature documenting that 1) copper can be mobilized in calcium-like waves upon certain stimuli^71^; 2) calcium mobilization itself can drive copper mobilization^95^; and 3) copper is mobilized during starvation stress to affect the enzymatic activity of Ulk1/2^71^,: we propose that this observation supports a parallel model where canonical calcium mobilization during ER stress induction results in copper mobilization to increase PERK occupancy and/or activity as an additional regulatory mechanism to finely tune PERK signaling in response to the stress environment. Thus, an increase in cytosolic copper after stress induction could mediate persistent PERK induction during long-term stress, while simultaneously inducing a counterbalancing negative feedback loop in typical PERK fashion, resulting in CTR1 downregulation in a copper-dependent fashion as previously established^96^. This may be partially mediated by PERK-dependent translational attenuation, although the effect appears more targeted as it is much more apparent for CTR1 than other monitored proteins, such as PERK, eIF2α, and actin. The ongoing development of probes for live-cell imaging of endogenous labile copper should allow the validation of these hypotheses in future studies^97,98^.

Finally, in Figure 5, we utilized the *C. elegans* model to confirm the *in vivo* translation of the regulation of PERK by copper, as dissected at the molecular and cellular level above, and to investigate the pursuant *in vivo* implications for copper homeostasis as a mediator of ER stress tolerance at the organism level. As predicted, the CBM animals recapitulated phenotypes previously documented in *pek-1* KO animals, namely reliance on *ire-1* for development and greater ER stress sensitivity. These findings not only demonstrate that PERK-copper binding is conserved through metazoans but further suggests the preservation of the exact kinase-dead phenotype when copper coordination is disrupted, although further confirmatory studies are required. The CBM animals were expected to mimic *pek-1* KO animals on the basis of a kinase-dead phenotype, if preserved with the corresponding mutations in the *C. elegans* model. In contrast, we considered that partial PERK activity, while significantly less *in vitro*, would be sufficient and not impact signaling and/or PERK-dependent outcomes at the whole organism level. Thus, we hypothesized that the singular 991A mutant, would result in an intermediate phenotype, if not the lack of a phenotype. Instead, we found an intermediate phenotype that was nearly, to just as dramatic as the CBM, demonstrating that the perturbation of PERK-copper regulation, even without complete kinase inactivation, translates to clear pathologic phenotypes at the organism level, thus highlighting the physiologic importance of this regulatory paradigm.

While the findings presented here are novel, retrospective analysis of the literature provides a surprising amount of support for copper regulation of PERK and its implications. Most notably, UPR induction, specifically PERK signaling, was recently documented in a mouse model of Wilson’s disease where copper is in excess^99^. Furthermore, PERK signaling has been shown to be cytotoxic in this context, contributing to cerebrovascular injury and apoptosis, which are alleviated by treatment with the copper chelator D-penicillamine, in combination with the traditional Chinese medicine GanDouLing. These findings support the notion that altered copper homeostasis can result in aberrant PERK signaling. In the same vein, another study documented that pyrrolidine dithiocarbamate-complexed copper exposure induced apoptosis in lung epithelial cells via ER stress, as well as mitochondrial stress^100^. In another study, drug-resistant renal cell carcinoma cells were shown to be effectively targeted for apoptosis, again via ER stress and mitochondrial pathways, following exposure to cuprous oxide nanoparticles (CONPs), without systematic toxicity^101^. Notably, the study highlighted apoptosis induction by ER stress-mediated mechanisms, with observations of strong UPR induction and ER swelling. The study also noted the accumulation of copper in the cytosol and the downregulation of copper regulatory chaperones upon CONP treatment, with many of the effects appearing to be mediated by such downregulation^101^. Induction of the cytotoxic UPR by both CONPs and the knockout of CCS and ATOX1 suggest not only the possibility of copper-mediated PERK activity but also a potential role for copper loading of PERK under basal settings.

The utilization of calcium by many chaperones as well as PERK’s identity as an ER calcium sensor and interplay with calcium signaling have resulted in the recognition of calcium’s roles in proteostasis and fine-tuning of stress responses during protein dyshomeostasis, which have been covered in a number of reviews^102,103^. Identification of copper regulation of PERK raises the possibility that copper also has a larger role in protein homeostasis, a hypothesis supported by other recently characterized copper-binding enzymes. A recent preprint reported the copper-binding activity of E2D clade of ubiquitin conjugases^73^. In that study, copper was shown not only to be involved in E2D activity but also to act as an allosteric modulator. Specifically, these enzymes, which facilitate the ubiquitination of proteins for digestion by the proteasome, were potently activated by copper supplementation in cell culture, leading to increased protein degradation. The study highlights p53 as a major target of these pathway and demonstrates that such signaling is required for morphogenesis, as flies harboring the mutants of the copper-binding sites of these enzymes were embryonically lethal when ubiquitously expressed and had major developmental issues when knocked in later. These recent discoveries on copper regulation of enzymes involved in signaling related to protein homeostasis, namely the unfolded protein response (PERK), autophagy (ULK), and proteasome activity (E2D ubiquitin ligase), invite an exciting new overarching signaling paradigm where copper, in concert with PERK and calcium, may act as a dynamic master regulator of protein homeostasis.

In closing, copper regulation of PERK, as shown here, paves the way for many future studies to thoroughly delineate the biochemical and cellular mechanisms underlying this regulation. Furthermore, recognition of this novel regulatory paradigm both explains and further implicates crosstalk of ER stress and other signaling responses in the determination of cell fate outcomes.

## Acknowledgements

This work was funded by NIH/NIMH R01MH109382 (K.J.S.), NIH T32GM008275 Structural Biology and Molecular Biophysics Training Grant (S.E.B.N), and the University of Pennsylvania BGS Training Grant (S.E.B.N.). Special thanks to the Koumenis Lab at the University of Pennsylvania for providing WT and KO PERK MEFs, the Brady lab at the University of Pennsylvania for providing CTR1, KO CTR1, ATP7A+/+ and ATP7A−/− MEFs, and the Gidalevitz lab at Drexel University and InVivo Biosystems for providing the *C. elegans* strains. Sequencing was completed by the Department of Genetics at the University of Pennsylvania.

## Author Contributions

Conceptualization, S.E.B.N, D.C.B, and K.J.S.; Methodology, S.E.B.N; Investigation, S.E.B.N., N.R.B, M.K.B., Validation, N.R.B, M.K.B, J.P. X.S.; Writing – Original Draft, S.E.B.N.; Writing –Review & Editing, S.E.B.N, C.A.E, J.P. T.G., D.C.B., K.J.S.; Funding Acquisition, K.J.S.; Resources, K.J.S., D.C.B and T.G.; Supervision, C.A.E., T.G., D.C.B, K.J.S.

## STAR Methods

### Resource Availability

#### Lead contact

Further information and requests for resources and reagents should be directed to and will be fulfilled by the lead contact, Kelly Jordan-Sciutto.

#### Materials availability

Plasmids generated for these investigation, as well as sequence files, are available upon request from the lead contact. The mutant *C. elegans* strains created by InVivo BioSystems are also available upon request

#### Data code and availability

All data reported in this paper (i.e. original western blot images, raw ELISA values, original *C. elegans* counts) will be shared by the lead contact upon request. This paper does not report original code. Any additional information required to reanalyze the data reported in this paper is available from the lead contact upon request.

### Experimental model and study participant details

#### Cell culture

WT and KO PERK MEFs, immortalized with SV40, were the kind gift of Dr. Koumenis^104^. CTR1 and KO CTR1, and ATP7A +/+ and −/− MEF lines were the generous gift of Dr. Donita Brady^67,71,105^. MEFs were cultured in DMEM with GlutaMAX and 10% FBS. According to experiment cells were treated with Thapsigargin (Tocris), PERK activator (Millipore), tunicamycin (Torcis), Cu-ATSM (Cayman Chemical), salubrinal (Sigma), TTM (Sigma), and BCS (Sigma) at various time points and doses as indicated in the results section. Whole-cell lysates were prepared utilizing NP-40 buffer (50 mM Tris pH 7.5, 120 mM NaCl, 0.5% NP-40, 0.4 mM Na_3_VO_4_, and Halt™ Protease and Phosphatase Inhibitor Cocktail), and protein was quantified by Bradford Assay using Bio-Rad reagent.

#### C. elegans

N2AM (wild-type) animals were a subclone of N2Bristol from the Morimoto lab provided by the Gidalevitz lab. *pek-1* null animals, strain TG157 (*pek-1(ok275); zcls4[phsp-4::GFP] +/-*), were previously generated by the Gidalevitz lab by crossing the following strains from the Caenorhabditis Genetic Center (CGC): SJ4005(*zcls4[phsp-4::GFP]V*) and RB545(*pek-1(ok275)X*)^87^. The mutant animals, strains COP2500(*pek-1(knu1130[M991A])*) and COP2509(*pek-1(knu1138[H931A, M99A1])*) were generated by InVivo Biosystems.

OP50 from the CGC and OP50 RNAi clone strains from the Ahringer library and supplied by the Gidalevitz lab at Drexel University^106,107^.

### Method Details

#### Cloning

PERK cDNA constructs were cloned from pTCN background gift constructs from Gerard Schellenberg^108^. The full length cDNA and kinase domain sequences were subcloned into a pcDNA3.1 C-terminal FLAG-tagged vector^109^. HRI cDNA in the pMD18 vector background was purchased from SinoBiological and subcloned into the same. Cloning was done by Gibson assembly utilizing NEB HiFi Assembly Master Mix. Mutations of the copper binding site were inserted using the Agilent Site Directed Mutagenesis kit. All construct sequences were verified with sequencing by the University of Pennsylvania, Department of Genetics Sequencing Core.

#### DNA Purification

DNA was produced by transformation of OneShot TOP10 chemically competent *E. coli* according by manufacturer instructions and grown in Luria Broth with Carbenicllin. DNA was purified using the ThermoFisher HiPure Purification kit according to manufacturer instructions. DNA concentrations were quantified on a Nanodrop 2000c.

#### Protein Expression and Purification

The protocol for protein expression and was derived from the protocol published by Portolano *et al.* generating a novel HEK 293F expression system^110^. Protein construct expression was performed by transfection into ThermoFisher FreeStyle 293F cells according to the expression system product manual, utilizing FreeStyle MAX transfection reagent diluted in OptiPRO SFM and 312.5ug of DNA per 250mL of culture. Expression proceeded for 2-3 days before harvesting by centrifugation for 5 minutes at 3,000g at 4°C. The pellet was resuspended with 1mL lysis buffer/25mL culture, composed of 25mM Tris-HCl, 150mM NaCl, 1mM EDTA, 1% NP-40, and 5% glycerol supplemented with Halt Phosphatase and Protease Cocktail Inhibitor Cocktail, mammalian protease inhibitor cocktail without chelators, and sodium orthovanadate, added immediately before use. Complete lysis was ensured by vortexing, homogenization, and sonication for 15 seconds, followed by 15 seconds of rest for 3 cycles, all on ice. Lysates were centrifuged for 25 minutes at 30,000g, 4°C to pellet cellular debris. The supernatant was collected and added to a spin column containing 1mL packed, rinsed, and equilibrated anti-FLAG resin from ThermoFisher. The lysate was incubated with the resin with agitation for 2 hours at 4°C. The column was then centrifuged 2 minutes at 1000g, 4°C, and the full-through collected for purification analysis. The resin was washed 4 times with 5 mL buffer, first with lysis buffer, then with lysis buffer + 300mM NaCl, incubated for 10 minutes at 4°C with agitation, twice, then with lysis buffer again. Washes were incubated for 10 min at 4°C with agitation. The protein was eluted using 2mL 1.25mg/mL 3X Flag Peptide from ThermoFisher in lysis buffer. Elutant was cleaned up by and concentrated using Amicon 10kda MWCO ultrafiltration columns, and repurified with anti-FLAG resin as necessary. Protein concentration measured was by Bradford Assay. Purity of final product was assessed by western blot and Coomassie staining.

#### IMAC pull-down

The protocols for copper-binding assays followed previously published metal pull-down experiments.^71,75^ Briefly, Bio-Rad IMAC resin was rinsed and charged with 200mM of different metal ions in 25mM Tris-HCL. Excess metal was then removed by 5 washes, and the charged resin incubated with cell lysate or purified protein for 2 hours at 4°C. Flow through was collected and bound protein was eluted with titrations of binding buffer (25mM Tris HCL, 100mM NaCL) plus varying concentrations of imidazole. Following washes/elution, sample buffer was added to the resin and SDS–PAGE analysis and immunoblotting were performed as described on all fractions.

#### Glutathione Pull-down

The protocols for copper-affinity assays followed previously published metal pull-down experiments^67,111^. For glutathione pull-down experiments GE glutathione sepharose was rinsed and charged with 50mM copper or left uncharged in protein storage buffer (1mM Tris-HCl, 150nM NaCl, 0.02% Triton-X, and 5% glycerol). Excess copper was removed by several washes and the resin was incubated with purified protein for 1-4 hours at 4°C. Resin was washed, with and without competitive copper or glutathione-complexed copper titrations. Following washes/elution, sample buffer was added to the resin and SDS–PAGE analysis and immunoblotting were performed as described on all fractions.

#### Kinase Assay

The *in vitro* kinase assay was derived from previous publications^67,75,112^. Kinase assays were prepared utilizing CST Kinase Buffer and 100µM ATP. For kinase assays with GST-tagged PERK Kinase domain, 20ng GST-PERK was combined with 7.3ng His-eIF2α in a 50uL reaction for a 4.2nM:4.2nM substrate:enzyme molar ratio. For kinase assays with FLAG tagged full-length PERK, 100ng was combined with 7.3ng His-eIF2α in a 50uL reaction for a 4.2nM:16.8nM molar ratio. After incubation, 50uL reactions were terminated with the addition of 4X NuPAGE LDS, 10X NuPAGE Sample Reducing Agent, and water up to 80uL. 20uL of the final reaction mixture was used for blotting. For kinase reactions with FLAG-tagged HRI, 630ng was combined with 500ng His-eIF2α in a 10uL reaction for a 1.4µM:875nM molar ratio. After incubation these reactions were terminated with 4X LDS, 10X Reducing Buffer, and H2O up to 270uL. 1uL of this solution was used for preparation of western blot samples, made up to 20uL with 4X LDS, 10X Reducing Buffer, and H2O.

#### ELISA

Kinase reactions for analysis by ELISA were prepared in 100uL reactions containing the same concentrations of PERK, eIF2α, buffer and ATP as above. Reactions were incubated at 37°C and terminated at the indicated time points by the addition of 21µM EDTA and transfer to ice. Reactions was diluted 1:1, processed using the Cell Signaling Technologies (CST) Pathscan p-eIF2α ELISA according to manufacturer instructions, and quantified using a ThermoFisher Multiskan™ FC Microplate Photometer at 450nm.

#### Gel Electrophoresis

Samples were prepped by combination with 4X LDS (Invitrogen) and 10X reducing buffer (Invitrogen) and heated for 10 min at 70°C. Samples were loaded into 4-12% Bis-Tris gels (Invitrogen) and ran at 130V in MOPS running buffer.

#### Western Blot

Gels for western blotting were transferred to PVDF membrane for 2.5 hours to overnight at 4°C in ThermoFisher Transfer buffer at 30V. Blots were then rinsed with Tris-buffered saline (TBS) and TBS with 0.1% Tween-20 (TBS-t) before being blocked 1 hour at room temperature in TBS-T + 5% BSA. Blot were incubated in primary antibody diluted in blocking buffer overnight. Primary antibodies and dilutions used were: 1:1000 Rabbit anti Ser 51 peIF2α (CST), 1:1000 Rabbit anti D9G8 peIF2α (CST), 1:1000 Mouse anti eIF2α (CST), 1:1000 Rabbit anti PERK (CST), 1:1000 Rabbit anti FLAG (CST), 1:40,0000 Mouse anti B-Actin (CST), 1:1000 Mouse anti CHOP (CST), 1:1000 Rabbit anti Cleaved Caspase 3 (CST), 1:1000 Rabbit anti CTR1 from (CST), 1:1000 Rabbit anti CCS from (Abcam), 1:1000 Rabbit anti ATP7A (Sigma), and 1:2000 Mouse anti BiP (BD). Blot were washed with TBS-T and incubated with 1:2000 Goat anti Rabbit or 1:5000 Goat anti Mouse HRP conjugated antibodies (Jackson) diluted in blocking buffer for 30 minutes at room temperature. Blots were washed an additional three time with TBS-T and one time with TBS before 3 minute exposure with ECL HRP substrate (Millipore) and imaged on a Bio-Rad ChemiDoc system. Images were quantified using ImageLab software.

#### LDH Assay

Roche LDH Cytotoxicity Assay was optimized and performed according to kit manufacturer instructions (Sigma). LDH activity was quantified using a ThermoFisher Multiskan™ FC Microplate Photometer at 492nm.

#### C. elegans

Standard methods were used for worm culture and genetics^113^. Briefly, *C. elegans* strains were maintained at 20°C on NGM media seeded with OP50. Strains were confirmed by PCR and sequencing. Animals were synchronized by picking gastrula-stage embryos, or by hypochlorite treatment.

The protocol for stress tolerance assays was derived from previously published results^89^. Specifically, drug treatments were added to fresh plates immediately before the beginning of all experiments. 10-30 gravid adults were allowed to lay eggs for approximately 4 hours and then 25-100 eggs per line per condition were moved to drug plates. Worm were counted and development monitored on days 2 and 3. The experiment was repeated 3 times with different populations of animals.

The protocols used for RNAi gene-specific knockdown and development assays were previously published^87^. For RNAi experiments, animals were grown for two generations on 0.4mM IPTG–containing plates, spotted with designated RNAi bacteria, either the L440 empty vector or *ire-1* RNAi. The RNAi plates were tested for the expected phenotypes, such as larval arrest of second generation of *pek-1* mutant animals on *ire-1* RNAi, to ensure proper induction of the RNAi. Specifically, gravid adults, grown from eggs on RNAi bacteria were allowed to lay eggs on control plates for approximately 4 hours. 20-50 of these second generation eggs were transferred to the corresponding RNAi plates and the development of these second generation animals was monitored 3 days later.

### Quantification and Statistical Analysis

All statistical analyses were performed using GraphPAD PRISM software. See figure legends for details of individual analyses.

## Supplemental Figure Titles and Legends

**Supplemental Figure 1-Selective pull-down of PERK by copper-charged resins**

**A)** Western blot images, probed for PERK, of 500ng purified GST-PERK kinase domain protein pulled-down by IMAC resin, charged as indicated, and eluted over an increasing imidazole titration. FT=Flow-through; Imidazole=20-100mM.

**B)** Western blot images, probed for PERK, of 500ng of purified PERK-FLAG kinase domain protein pulled-down by glutathione sepharose, charged as indicated.

**C)** Western blot images, probed for PERK, of 100ng of purified full-length PERK-FLAG protein pulled-down by glutathione sepharose, charged as indicated. Wash=binding buffer plus+50mM copper sulfate.

**Supplemental Figure 2-Kinase assay conditions**

**A)** Plot of enzyme progress curve (p-eIF2α accumulation as monitored by absorbance at 450nm, normalized to PERK, over time) for ELISA analyzed kinase assay reactions at 37°C Data was fit by least squares regression to an exponential plateau model. With Y_0_ constrained to 0, Y_m_ constrained to less than 4 (the detection limit on the plate reader), and k constrained to greater than 0, the exact sum of squares F test was used to determine if the best-fit values of unshared parameters differed between the curves fitted to each construct, as compared to a global model; p=0.0304.

**B)** Table of parameters for curves fit in A. Notably, 95% confidence intervals could not be calculated for the curve parameters.

**C)** Coomassie images of purified full-length PERK protein preparations used to prepare ELISAs. Relative densitometry quantifications below used for normalization in A.

**D,F,H)** Western blot images, probed as indicated, of full-length PERK-FLAG(D), HRI-FLAG(F), or GST-PERK kinase domain(H) kinase assay reactions carried out at 4°C(D), 37°C(F), or 25°C(H) and terminated at the indicated time points.

**E,G,I)** Quantification of the experiments in D(E), F(G), or H(I) plotted as enzyme progress curves. Data was fit by least squares regression to an exponential plateau model with Y_0_ constrained to 0, and Y_m_ and k constrained to greater than 0. R^2^=0.9683(E), R^2^=0.6028(G), or R^2^=0.8991(I). Runs test gave non-significant deviation from the model with p=0.9714(E), p=0.8(G), or p=0.8(I).

**Supplemental Figure 3-Evidence of copper-dependent regulation of PERK-dependent eIF2α phosphorylation in the cell**

**A,C)** Representative western blot images, probed as indicated, of WT and KO PERK (gray) MEF whole cell lysates from cells treated with 15um TTM for 24 hours, and 300nM Tg for 2 hours(A) or 50µM Sal for 24 hours(C).

**B,D)** Quantification of n=3-5 replicates of the experiments in A(B) or C(D). One-way repeated measures ANOVA returned no significant effect for Sal treatment (D, p=0.131) or Tg treatment for KO PERK (B, gray squares, p=0.185) but returned a significant effect for WT (p<0.0001), with Holm-Sidak’s multiple comparison tests returning significance for each group in comparison to Tg+Veh: *p<0.05; **p<0.01.

All quantifications plot individual n as data points and each bar represents mean±SD. Data is normalized by experimental replicate as fold changes from p-eIF2α induction alone(Tg or Sal) as indicated by the dotted line at y=1.

**Supplemental Figure 4-Copper homeostasis and ER stress tolerance**

**A,B)** Quantification of untreated p-eIF2α from western blot images of four replicates of the experiment shown in Figure 4 A(A) or F(B). Data is normalized as the fold change from the average untreated ratio, pooled across all samples. Two-tailed unpaired t-tests: **p<0.01.

**C,D)** LDH Assay results quantifying LDH release as a measure of cytotoxicity from three replicates each of WT and KO PERK (gray squares) MEFs treated with 300nM Tg(C) or 0.5ug/mL Tm(D) for 20 hours. Cytotoxicity is calculated as a percentage of the total LDH released by lysed cells, after normalization by subtracting out any LDH release signal from untreated cells. Two-tailed paired t-tests: *p<0.05.

**E)** Western blot images, probed as indicated, of WT and KO PERK (gray) MEF whole cell lysates from cells treated for with 300nM Tg or 0.5ug/mL Tm for 20 hours.

All quantifications plot individual n as data points (n=3-4) and each bar represents mean±SD.

**Supplemental Figure 5-Evidence of altered ER stress tolerance in PERK copper-binding mutant animals**

**A, B)** Development/Survival analysis of *C. elegans* eggs of each strain incubated on 1ug/mL Tm for 3 days. Chi-squared=42.30, p<0.0001(A), or 9.617, p=0.0571(B). Biological replicates of Figure 5B, included in 5C.

**C)** Development/Survival analysis of *C. elegans* eggs of each strain incubated on DMSO vehicle for 1ug/mL Tm for 3 days.

