## Supplemental Data for "PERK Kinase Activity is Regulated by Copper Binding-A New Regulatory Paradigm for Modulation of ER Stress Tolerance"

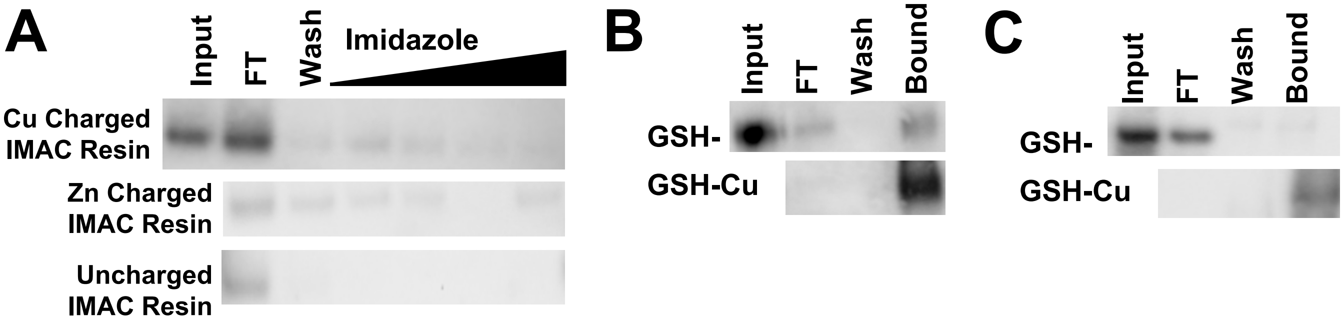


***Supplemental Figure 1-Selective pull-down of PERK by copper-charged resins***

**A)** Western blot images, probed for PERK, of 500ng purified GST-PERK kinase domain protein pulled-down by IMAC resin, charged as indicated, and eluted over an increasing imidazole titration. FT=Flow-through; Imidazole=20-100mM.

**B)** Western blot images, probed for PERK, of 500ng of purified PERK-FLAG kinase domain protein pulled-down by glutathione sepharose, charged as indicated.

**C)** Western blot images, probed for PERK, of 100ng of purified full-length PERK-FLAG protein pulled-down by glutathione sepharose, charged as indicated. Wash=binding buffer plus+50mM copper sulfate.

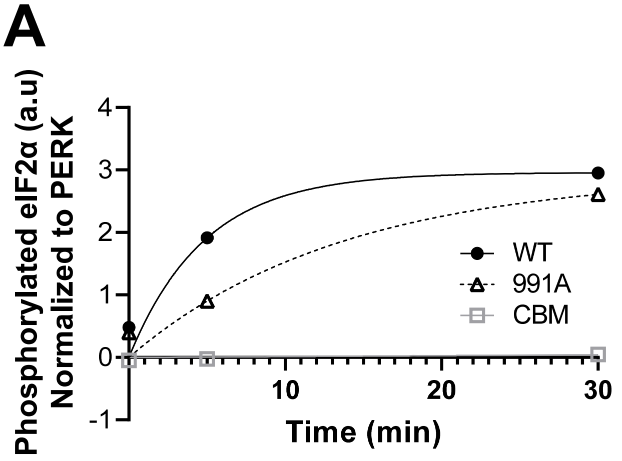


B

| | Ym | k | R2 | Runs |
|---|---|---|---|---|
| WT | 2.959 | 0.207<br>8 | 0.924<br>1 | n.s. |
| 991<br>A | 2.932 | 0.073<br>5 | 0.940<br>5 | n.s.<br>p>0.99 |
| CBM | 4 | 0.000<br>3 | 0.344<br>1 | n.s.<br>p=0.67 |

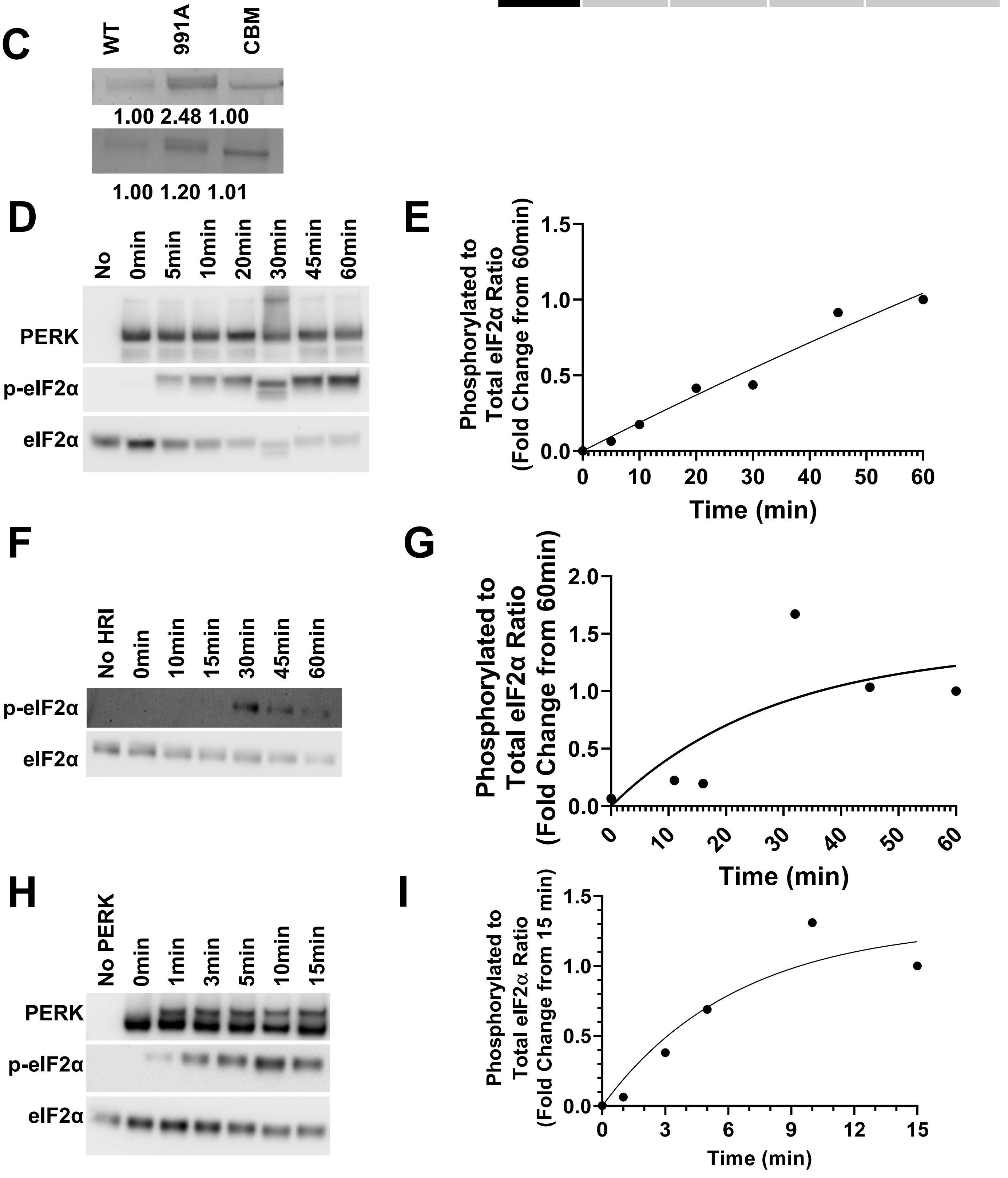

***Supplemental Figure 2-Kinase assay conditions***

**A)** Plot of enzyme progress curve (p-eIF2α accumulation as monitored by absorbance at 450nm, normalized to PERK, over time) for ELISA analyzed kinase assay reactions at 37°C. Data was fit by least squares regression to an exponential plateau model. With $Y_0$ constrained to 0, $Y_m$ constrained to less than 4 (the detection limit on the plate reader), and k constrained to greater than 0, the exact sum of squares F test was used to determine if the best-fit values of unshared parameters differed between the curves fitted to each construct, as compared to a global model; p=0.0304.
**B)** Table of parameters for curves fit in A. Notably, 95% confidence intervals could not be calculated for the curve parameters.
**C)** Coomassie images of purified full-length PERK protein preparations used to prepare ELISAs. Relative densitometry quantifications below used for normalization in A.
**D,F,H)** Western blot images, probed as indicated, of full-length PERK-FLAG(D), HRI-FLAG(F), or GST-PERK kinase domain(H) kinase assay reactions carried out at 4°C(D), 37°C(F), or 25°C(H) and terminated at the indicated time points.
**E,G,I)** Quantification of the experiments in D(E), F(G), or H(I) plotted as enzyme progress curves. Data was fit by least squares regression to an exponential plateau model with $Y_0$ constrained to 0, and $Y_m$ and k constrained to greater than 0. $R^2$=0.9683(E), $R^2$=0.6028(G), or $R^2$=0.8991(I). Runs test gave non-significant deviation from the model with p=0.9714(E), p=0.8(G), or p=0.8(I).

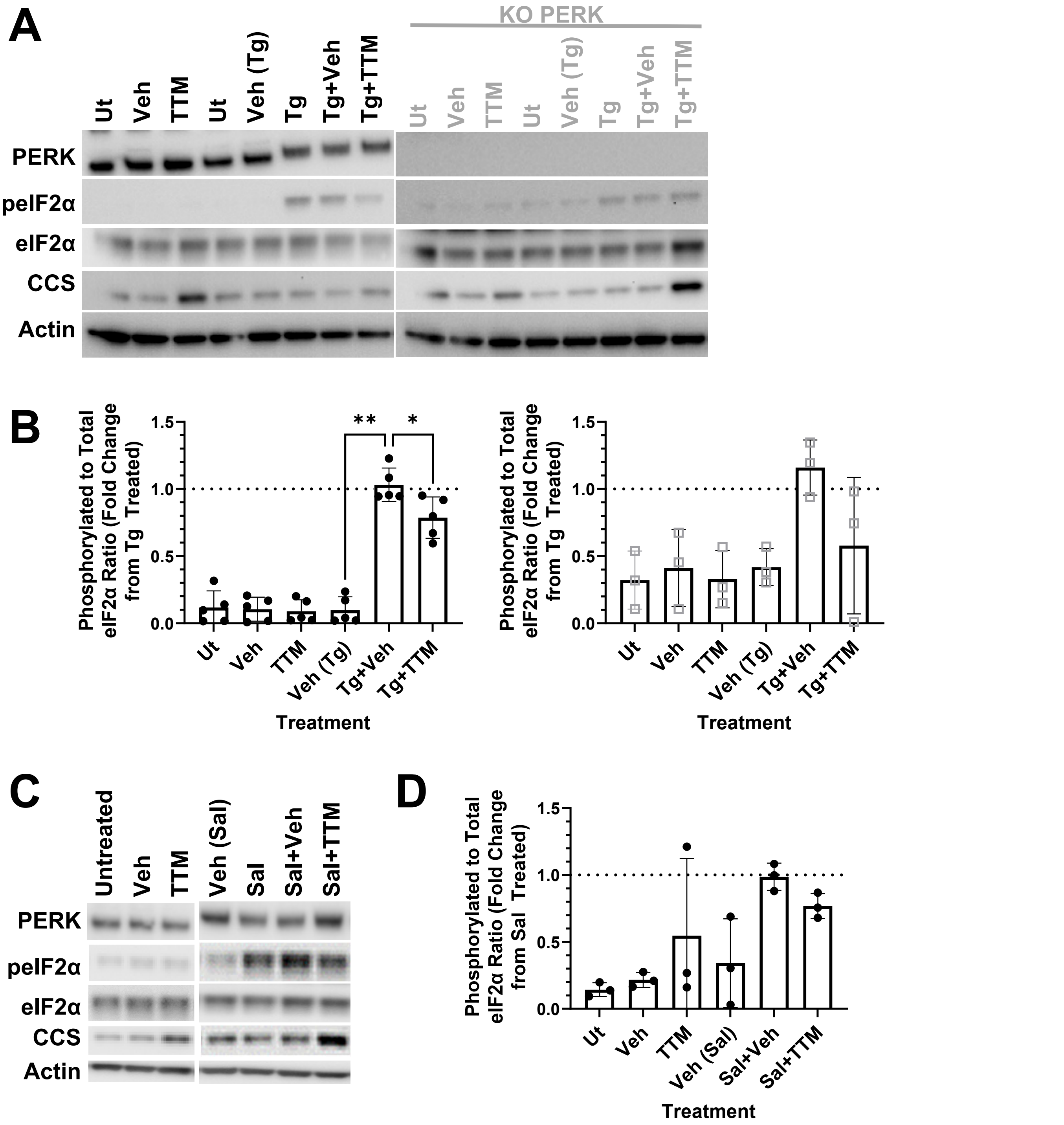


***Supplemental Figure 3-Evidence of copper-dependent regulation of PERK-dependent eIF2α phosphorylation in the cell***

**A,C)** Representative western blot images, probed as indicated, of WT and KO PERK (gray) MEF whole cell lysates from cells treated with 15um TTM for 24 hours, and 300nM Tg for 2 hours(A) or 50μM Sal for 24 hours(C).

**B,D)** Quantification of n=3-5 replicates of the experiments in A(B) or C(D). One-way repeated measures ANOVA returned no significant effect for Sal treatment (D, $p=0.131$) or Tg treatment of KO PERK MEFs (B, gray squares, $p=0.185$), but returned a significant effect for WT ($p<0.0001$), with Holm-Sidak's multiple comparison tests returning significance for each group in comparison to Tg+Veh: *$p<0.05$; **$p<0.01$.

All quantifications plot individual n as data points and each bar represents mean±SD. Data is normalized by experimental replicate as fold changes from p-eIF2α induction alone(Tg or Sal) as indicated by the dotted line at y=1.

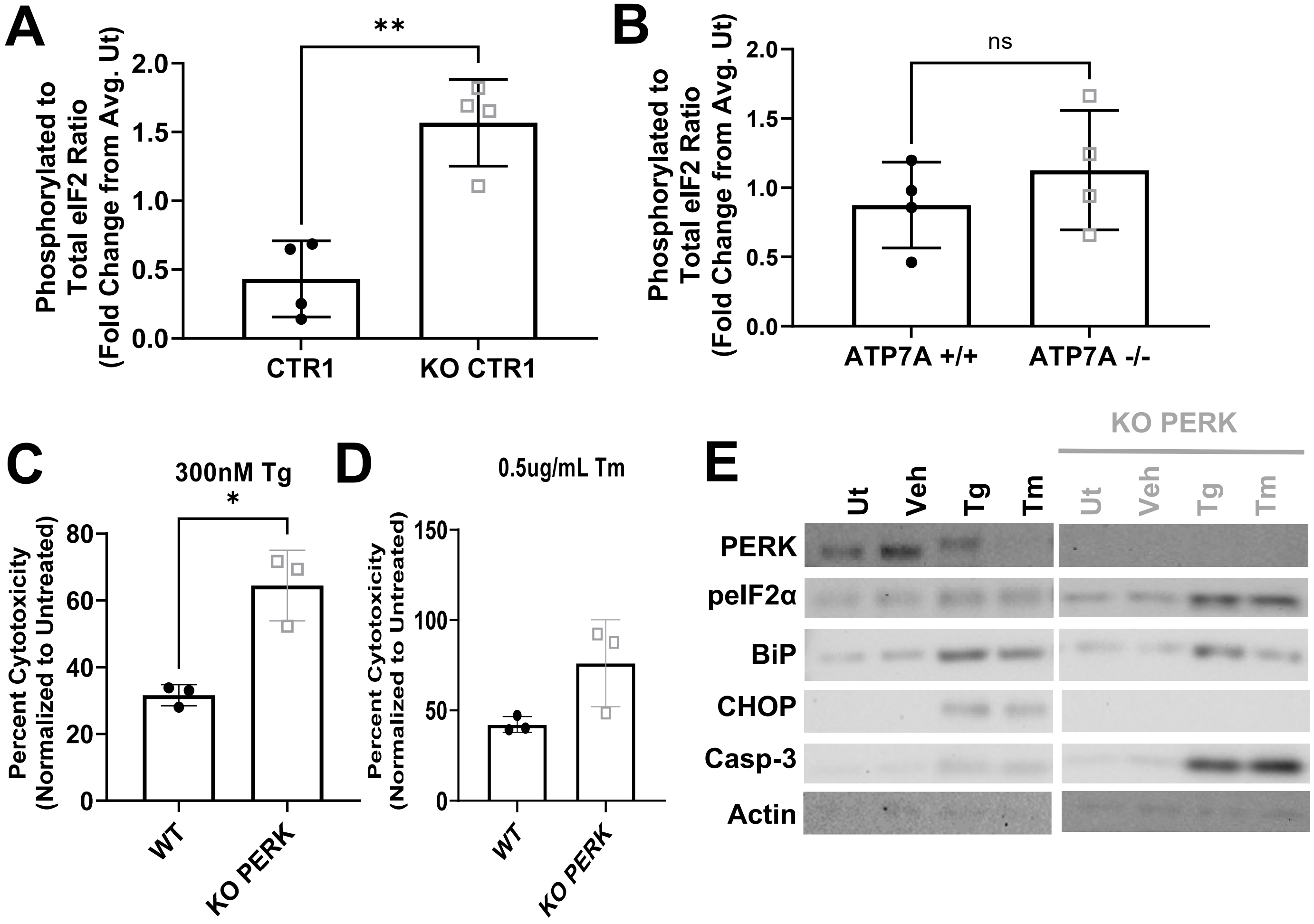


***Supplemental Figure 4-Copper homeostasis and ER stress tolerance***

**A,B)** Quantification of untreated p-eIF2α ratios from western blot images of four replicates of the experiment shown in Figure 4 A(A) or F(B). Data is normalized as the fold change from the average untreated ratio, pooled across all samples. Two-tailed unpaired t-tests: **p<0.01.

**C,D)** LDH Assay results quantifying LDH release as a measure of cytotoxicity for WT and KO PERK (gray squares) MEFs treated with 300nM Tg(C) or 0.5ug/mL Tm(D) for 20 hours. Cytotoxicity is calculated as a percentage of the total LDH released by lysed cells, after normalization by subtracting out any LDH release signal from untreated cells. Two-tailed paired t-tests: *p<0.05.

**E)** Western blot images, probed as indicated, of WT and KO PERK (gray) MEF whole cell lysates from cells treated for with 300nM Tg or 0.5ug/mL Tm for 20 hours.

All quantifications plot individual n as data points (n=3-4) and each bar represents mean±SD.

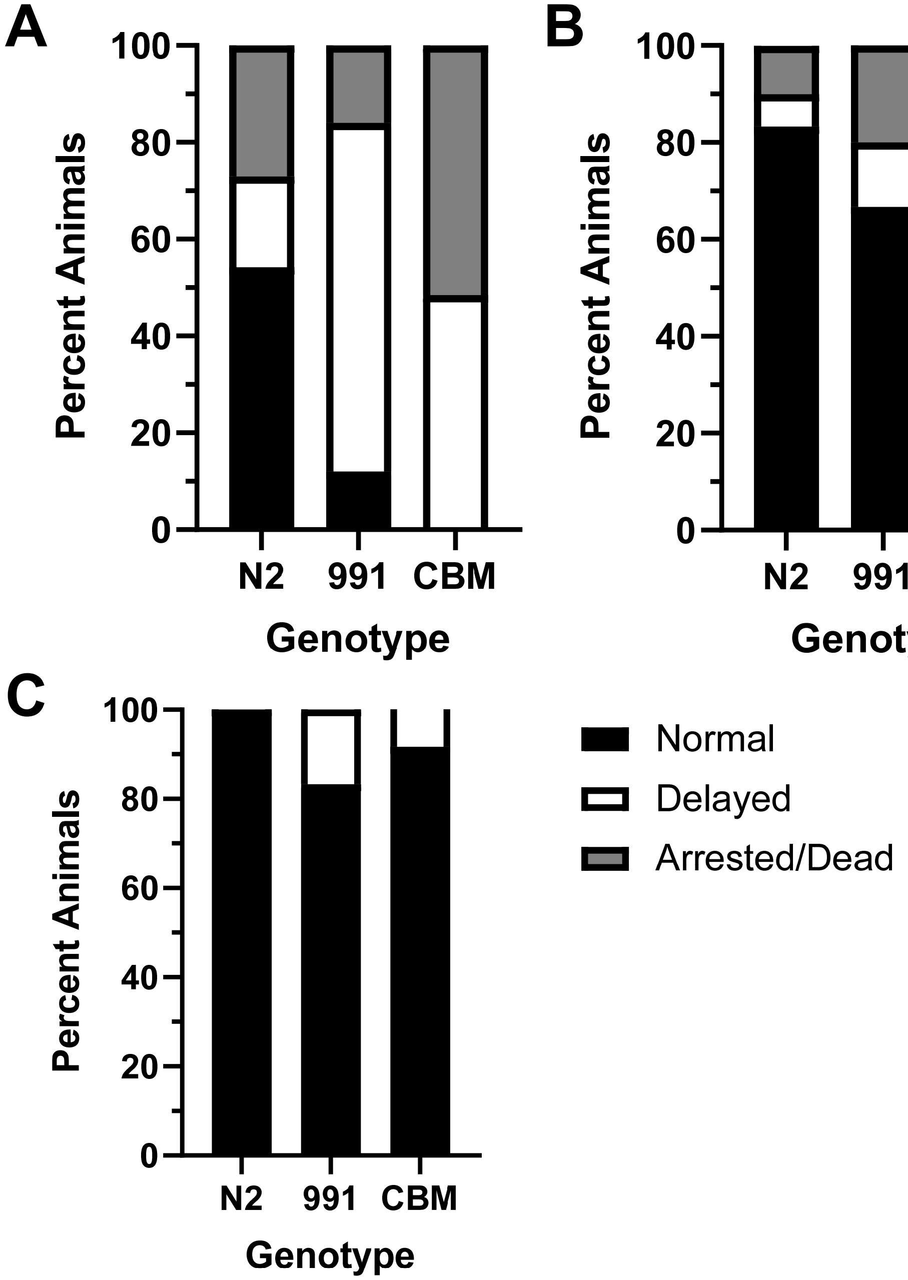


***Supplemental Figure 5-Evidence of altered ER stress tolerance in PERK copper-binding mutant animals***

**A,B)** Development/Survival analysis of *C. elegans* eggs of each strain incubated on 1ug/mL Tm for 3 days. Chi-squared=42.30, p<0.0001(A), or 9.617, p=0.0571 (B). Biological replicates of Figure 5B, included in 5C.

**C)** Development/Survival analysis of *C. elegans* eggs of each strain incubated on DMSO vehicle for 1ug/mL Tm for 3 days.
